# Differentiation of human T follicular helper cells *in vitro* requires co-operation between STAT3 and SMAD signaling cytokines that is restrained by IL-2

**DOI:** 10.1101/2025.01.12.632620

**Authors:** William S. Foster, Lucía Labeur-Iurman, Geoffrey S. Williams, Matthew J. Robinson, Gianluca Carlesso, Clare M. Lloyd, James A. Harker

## Abstract

T follicular helper cells (TFH) are a CD4 T cell subset required for effective humoral immunity; promoting antibody responses to vaccines and natural infections. The mechanisms by which human TFH differentiate are not fully understood, and given the anatomical localization to the secondary lymphoid tissues, and asynchronous nature of *in vivo* TFH populations, studying TFH differentiation *in vitro* represents an attractive alternative, through which the mechanisms which govern TFH differentiation can be explored. Here we demonstrate reliable differentiation of *in vitro* TFH from naïve human CD4 T cells in a STAT3 dependent fashion. Cooperation between STAT3 and SMAD signaling cytokines led to expression of the lineage defining transcriptional regulator BCL6, multiple downstream proteins required for TFH function and localization (including PD1 and CXCR5), and potent IL-21 production capacity by these *in vitro* TFH. Critically we found that autocrine production of IL-2 acted as an intrinsic brake on TFH differentiation *in vitro*; promoting proliferation but limiting the acquisition of a TFH phenotype resembling that found in germinal centers *in vivo*; neutralization of IL-2 resulting in the emergence of this phenotype. Overall, we show that human TFH expansion, differentiation and function relies on careful regulation of pro- and anti-TFH cytokine signaling.

## Introduction

During an immune response, naïve CD4 T cells differentiate into specialized subsets of CD4 T cells with distinct functions. These T helper (TH) subsets are defined by the upregulation of lineage defining master transcription factors; for example TH1, TH2 and TH17 cells require TBET, GATA3 and RORγT respectively, to produce their hallmark cytokines IFNγ/TNFα, IL-4/5/13 and IL-17 respectively (1).

A more recently identified CD4 T cell subset, Follicular helper T cells (TFH) specialize in regulating antibody mediated immunity. They achieve this by rapidly upregulating the chemokine receptor CXCR5 upon priming, which promotes their migration toward and into the B cell zone of secondary lymphoid tissue. Here they can form close interactions with activated B cells to promote B cell survival, proliferation and differentiation through the expression of co-stimulatory molecules such as CD40L and cytokines including IL-21 and IL-4 (2). Unlike other CD4 T cell subsets, TFH require expression of a transcriptional repressor, BCL6, for their differentiation, which is necessary and sufficient for TFH differentiation in mice (3–5). In humans however, full differentiation also requires expression of the transcription factor MAF (6). TFH differentiation in both mice and humans is antagonized by BLIMP-1 (*PRDM1*); a negative regulator of BCL6 (3).

CD4 T cell fate is heavily influenced by extracellular cues, and in particular the cytokine micro-environment. Naive CD4 T cells can be differentiated into TH1 cells by exposure to IL-12 when IL-4 is suppressed and further activated by IL-18 (7). Inversely IL-4 is sufficient to promote TH2 cell differentiation when IFN-γ is suppressed. Meanwhile TH17 cells differentiation utilizes cytokines such as IL-6 and IL-23 that signal through Signal transducer and activator of transcription (STAT)-3, in combination with TGF-β.

In mice, STAT3 is also essential for TFH differentiation and its genetic deletion prevents TFH differentiation *in vivo* (8). Indeed, *in vitro* the potent STAT3 activating cytokine IL-6 promotes IL-21 production and early Bcl6 upregulation in CD4 T cells (9, 10), but is redundant for TFH differentiation *in vivo* in a number of infection and vaccination settings (11, 12). IL-6 may however be necessary for their long-term maintenance (12, 13). Simultaneous removal of IL-6 alongside IL-21, which also signals primarily through STAT3, does however result in reduced TFH differentiation after acute viral infection (14). Fitting with the importance of STAT3 signaling cytokines in TFH differentiation and function gp130, the common signal transduction molecule of the IL-6 cytokine family, is required for optimal TFH differentiation in chronic viral infection and allergic airway disease (15, 16), with IL-27 thought to synergize with IL-6 in this process (16). TGF-β signaling has also been implicated in TFH, promoting their formation in the context of respiratory infection, but conversely inducing apoptosis of TFH and preventing their accumulation during systemic immunization and limiting their ability to produce IL-21 and ICOS (17, 18). Conversely STAT-5 signaling, especially when driven by IL-2, inhibits the differentiation of TFH (19, 20). Indeed, conditions where IL-2 signaling is enhanced, such as early life, show limited TFH differentiation(21). Murine models have also suggested a somewhat distinct, stepwise requirement for co-stimulation in the differentiation of TFH, with both ICOS and CD28 signaling required for both priming and maintenance of TFH (22, 23).

Human TFH appear to rely on similar, but distinct, signaling pathways to those observed in mice. CD4 T cells from patients with a functional deficiency in *STAT3* have a reduced frequency of CXCR5^+^ CD4 T cells and a reduced capacity to produce IL-21 and help B cells upon stimulation (24). Likewise, individuals with mutations in *CD40L* and *ICOS* are associated with reduced TFH formation and hypogammaglobinemia (25–27). While IL-6 can also promote initial BCL6 up-regulation and IL-21 production, IL-12, which in man signals through both STAT3 and STAT4, appears to have a more prominent role (28, 29) with mutations that lead to defective *IL12RB* signaling being associated with reduced TFH frequencies, particularly in early life (30). TGF-β1, and another SMAD-signaling cytokine Activin A (ActA), also promote acquisition of key TFH associated molecules including BCL6 and IL-21, a process enhanced by IL-1β (31, 32). Meanwhile low dose IL-2 treatment of patients with autoimmunity reduces circulating TFH frequencies, supporting its role as a negative regulator of TFH (33). Human TFH appear to display significant plasticity, expressing both BCL6 and transcription factors and functions associated with other CD4 lineages, resulting in subtypes (34). For example IL-12 signaling can result in TFH that co-express T-BET and IFN-γ (35), while exposure to IL-23, a potent STAT3 activating cytokine, upregulates both BCL6 and RORγt (36), whilst IL-13 and GATA3 expressing TFH can also be observed in certain TH2 environments (37).

The sequence of cues required to promote neutral TFH generation from naïve T cells is however unclear, and direct comparisons of the importance of each cytokine shown to support TFH differentiation, given their shared use of STAT and SMAD signaling intermediaries are lacking. Here we therefore utilized a combination of factors shown to regulate TFH differentiation. We show that a combination of cytokines signaling through both the STAT3 pathway, IL-6 and IL-12, and the SMAD pathway, TGF-β1 and Activin-A, are capable of promoting TFH-like differentiation of naïve CD4 T cells, with some, but not all, features of TFH *in vivo*. Indeed, autocrine production of IL-2 by *in vitro* TFH naturally limited TFH differentiation, and neutralization of IL-2 allowed further acquisition of TFH phenotype and function.

## Results

### TFH-like cells can be generated from naïve human CD4 T cells in vitro

To mimic conditions thought to favor TFH differentiation *in vivo,* naïve peripheral blood CD4 T cells were first exposed to anti-CD3 to stimulate the TCR in the presence of CD28 and ICOS-L co-stimulation. After 24 hours the cells were then cultured in the presence of IL-6, IL-12, IL-1β, TGF-β and Activin A, all cytokines reported to favor certain aspects of TFH (*in vitro* culture schematic shown in Fig 1A).

**Figure 1.**
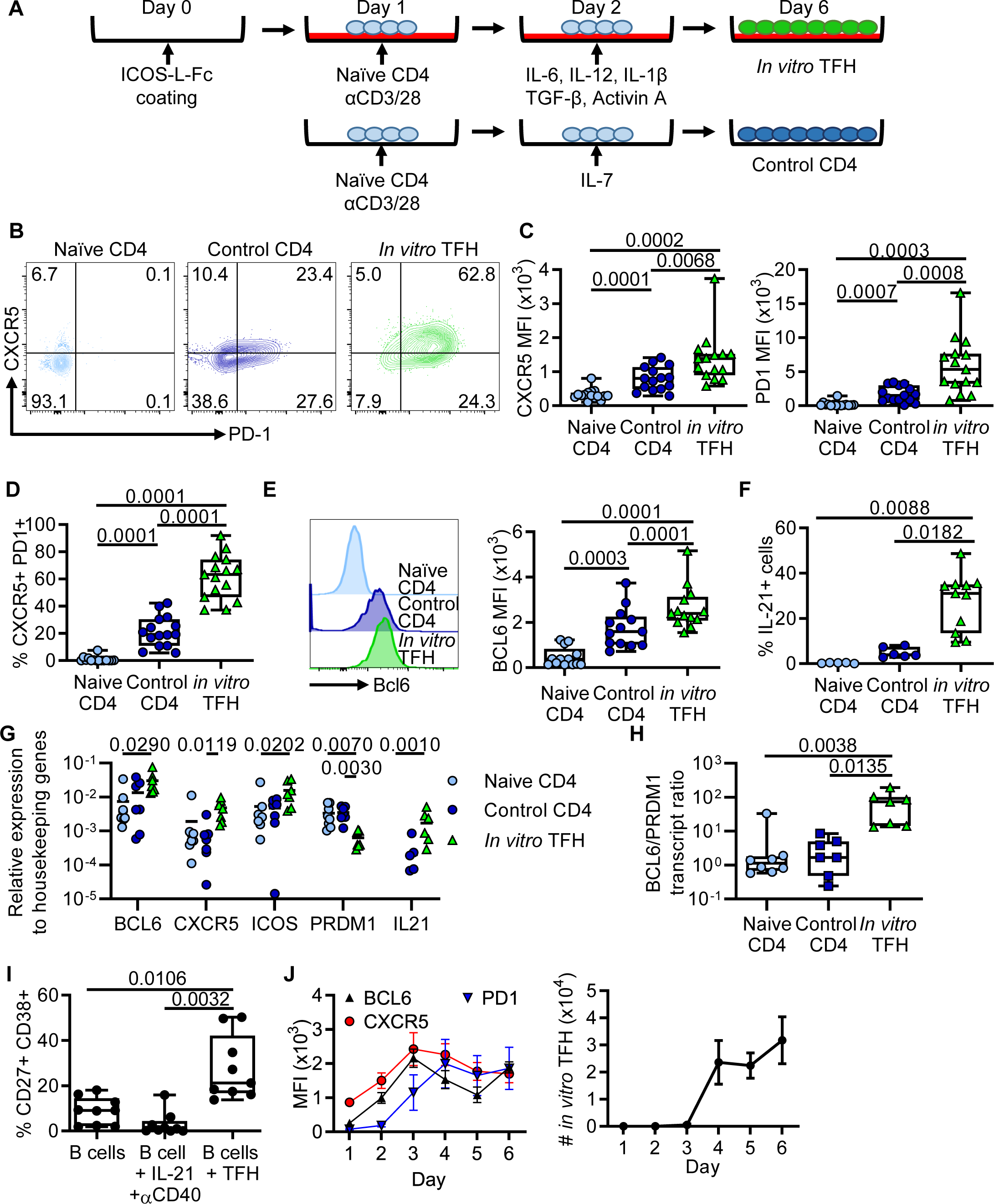
Human TFH analogs can be generated *in vitro*. **(A)** Naïve CD4 T cells were isolated from peripheral blood and cultured as displayed. Cells were harvested at day 6. **(B)** Representative plots of PD-1 and CXCR5 expression by naïve CD4 T cells, control CD4 and *in vitro* TFH cultured as in (A). Numbers represent percent of cells in each gate. **(C)** CXCR5 MFI, PD1 MFI and **(D)** proportion of CXCR5+ PD1+ cells generated (as gated in (B)). **(E)** Representative histogram and quantification of BCL6 expression. **(F)** IL-21 production was quantified after cells were stimulated with PMA/Ionomycin in the presence of Brefeldin A. **(G)** Expression of core TFH-associated transcripts was quantified, in relation to expression of housekeeping gene *ACTB* and *GAPDH* transcripts. **(H)** Ratio of BCL6:PRDM1 transcript was quantified from data in (G). **(I)** Heterologous naïve B cells were cultured with either 100ng/mL IL-21 and 5μg/mL αCD40 or *in vitro* TFH for 72 hours. **(J)** BCL6, CXCR5 and PD1 upregulation was quantified daily on a subset of *in vitro* TFH cultures. For **(C-I)** Tukey’s multiple comparison test was used.

These “TFH favoring” conditions led to significant and uniform upregulation of CXCR5 and PD1 at day 6 of culture compared to both the initial naïve CD4 T cells and CD4 T cells cultured under control conditions of anti-CD3, anti-CD28 and IL-7 (Fig 1B and C), resulting in a substantial population of CXCR5^+^ PD1^+^ *in vitro* TFH (Fig 1D). While BCL6 upregulation was seen after TCR stimulation in control cells, BCL6 expression was significantly upregulated in *in vitro* TFH compared to both naïve and control CD4 T cells (Fig 1E). *In vitro* TFH had greatly increased capacity for IL-21 production, following re-stimulation with PMA and Ionomycin (Figure 1F). These observations were confirmed at an RNA transcript level, with *in vitro* TFH having increased BCL6, CXCR5, ICOS and IL21 transcripts, along with decreased PRDM1 transcript as compared to naïve CD4 cells and control cultured CD4 cells (Fig 1G). Consequently, *in vitro* TFH had an increased ratio of BCL6:PRDM1 transcript (Fig 1H), further demonstrating their specific differentiation into the TFH lineage (3). Heterologous co-culture of *in vitro* TFH with naïve B cells resulted in the activation of B cells (Fig 1I). Analysis of the kinetics of TFH differentiation revealed CXCR5 and BCL6 were both rapidly upregulated, reaching maximal expression by day 3 of culture, while PD1 upregulated was slightly delayed with maximal expression achieved from day 4 onwards (Fig 1J). Expression of all 3 proteins was most stable at day 6 of culture, and the total number of CXCR5^+^ PD1^+^ cells generated continued to increase over time (Fig 1F). Overall, this data showed that by exposing naïve CD4 T cells to a specific cytokine and co-stimulatory environment *in vitro*, a homogenous population of CXCR5^+^ PD1^+^ CD4 T cells, with increased BCL6 expression and IL-21 production with B cell helper capacity can be generated.

### IL-12 and TGF-β play dominant and distinct roles in TFH differentiation in vitro

Next, the precise role of key mediators in driving *in vitro* TFH differentiation was determined. In humans IL-6 and IL-12 share common downstream signaling pathways including the phosphorylation of STAT3. Removal of both IL-12 and IL-6 from the *in vitro* culture resulted in a significant reduction in the surface expression of both CXCR5 and PD1, and a trend towards reduced BCL6 (Fig 2A). This was mirrored at a transcript level, and *IL21* expression was also found to be significantly lower (Fig 2B). Further, while BCL6 was not significantly down-regulated in the absence of IL-6 and IL-12, the ratio of *BCL6* to *PRDM1* (the gene that encodes BLIMP1) was significantly reduced. Removal of IL-6 alone appeared to have limited effects of TFH differentiation, while removal of IL-12 resulted in reductions in CXCR5 and *IL21*. Taken together this suggests IL-12 and IL-6 can act redundantly to promote TFH differentiation, especially upregulation of CXCR5 and IL-21, and promote an enhanced BCL6 to BLIMP1 ratio, with IL-12 playing the dominant role.

**Figure 2.**
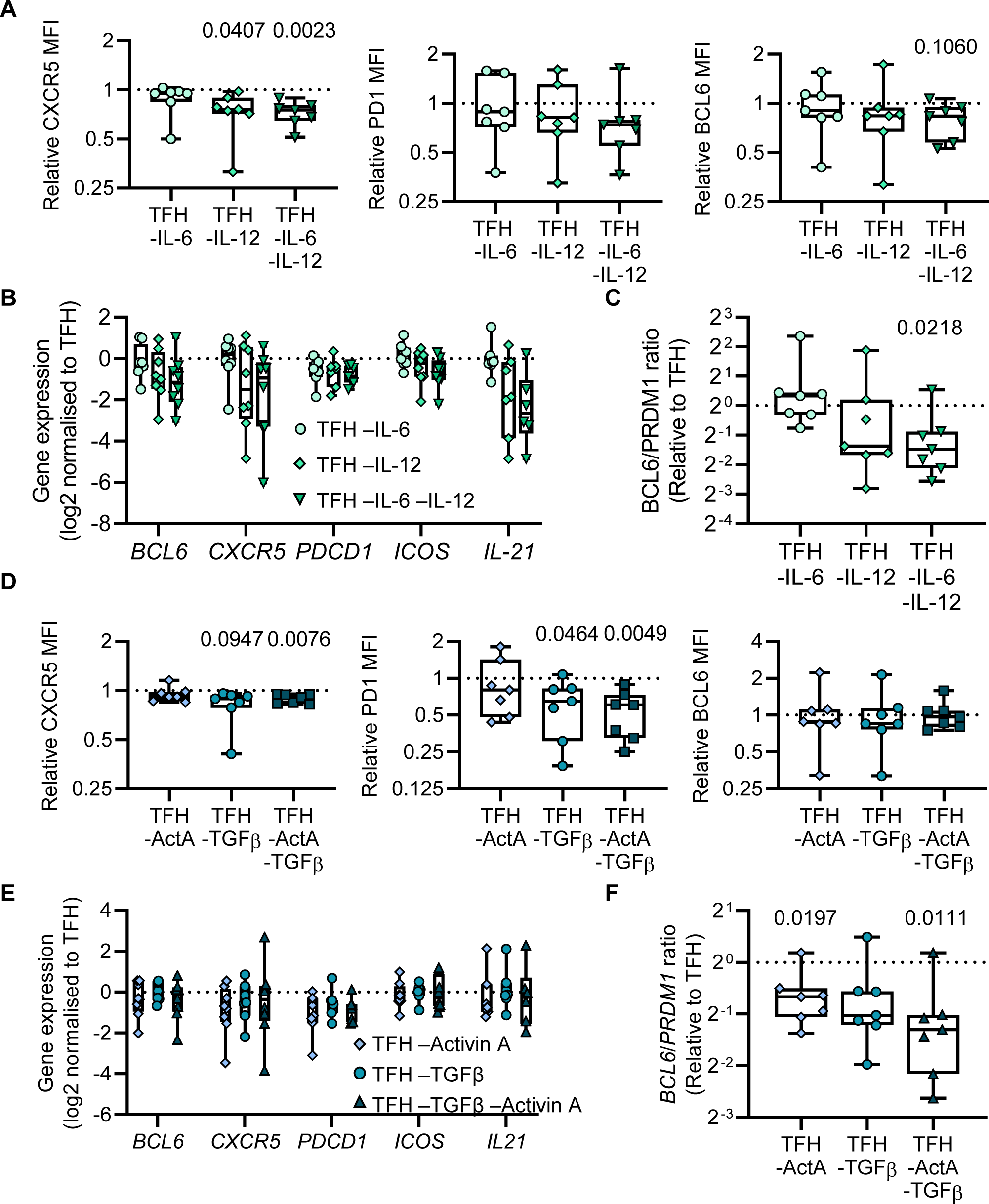
STAT and SMAD signaling cytokines promote specific TFH phenotypes. **(A-F)** *In vitro* TFH were generated as in Figure 1, with indicated cytokines removed. Data is presented as relative values as compared to matched *in vitro* TFH generated with a complete set of cytokines. **(A)** Relative CXCR5, PD1 and BCL6 MFI was quantified by flow cytometry. **(B)** Relative mRNA expression of *BCL6*, *CXCR5*, *PDCD1*, *ICOS* and *IL21.* **(C)** Ratio of *BCL6* and *PRDM1* mRNA. **(D)** Relative CXCR5, PD1 and BCL6 MFI was quantified by flow cytometry. **(E)** Relative mRNA expression of *BCL6*, *CXCR5*, *PDCD1*, *ICOS* and *IL21.* **(F)** Ratio of *BCL6* and *PRDM1* mRNA. For **(A-F)** n=7.

Unlike IL-12 and IL-6, TGF-β and Activin A both typically signal through the SMAD signaling cascade. Loss of TGF-β and Activin A resulted in downregulation of PD1 and to a lesser extent CXCR5 at the protein level (Fig 2D). There were no significant changes in the transcripts for *BCL6*, *CXCR5*, *PDCD1, ICOS* or *IL21* but due to an up-regulation of *PRDM1* the ratio of *BCL6* to *PRDM1* was reduced (Fig 2E & F). Loss of TGF-β alone resulted in similar reductions in CXCR5 and PD1, while loss of Activin A alone had no observable affect, suggesting a dominant role of TGF-β, however loss of both was required for changes in BCL6 to PRDM1 ratio to be observed.

Overall this data supports a role for cytokines signaling through both the SMAD and STAT pathways as playing distinct and overlapping roles in driving acquisition of a TFH-like phenotype, with SMAD signaling cytokines being particularly potent at promoting PD1 expression while STAT signaling cytokines were involved in CXCR5 and IL-21 upregulation.

### STAT3 is essential for TFH differentiation and function

Given the potent effects of IL-12, especially in combination with IL-6, during *in vitro* TFH differentiation the role of downstream signaling through either STAT3 or STAT4 was next investigated. Differentiation of naïve CD4 T cells into TFH in the presence of 100μM (±) Lisofylline, a small molecule inhibitor of IL-12 signaling and STAT4 activation, had little effect of acquisition of key TFH associated molecules including CXCR5, PD1 and BCL6 (Fig S1).

To inhibit STAT3 signaling, CD4 T cells were treated with C188-9 - an inhibitor of STAT3 phosphorylation, from 24 hours post culture. Strikingly treatment with 5 µM of C188-9, a concentration routinely shown to inhibit the majority of STAT3 phosphorylation(38), there was significant cell death at 6 days post culture in TFH differentiation, but not control conditions, when compared to vehicle treated cells (Fig 3A). Use of lower concentrations of C188-9 prevented this cell death, and there was limited effect on the expression of CXCR5, PD1 or BCL6 compared to vehicle treated cells (Fig 3B & C). Even at these lower treatment doses, there was however a significant reduction in the concentrations of CXCL13 and IL-21 in the supernatant (Fig. 3D).

**Figure 3.**
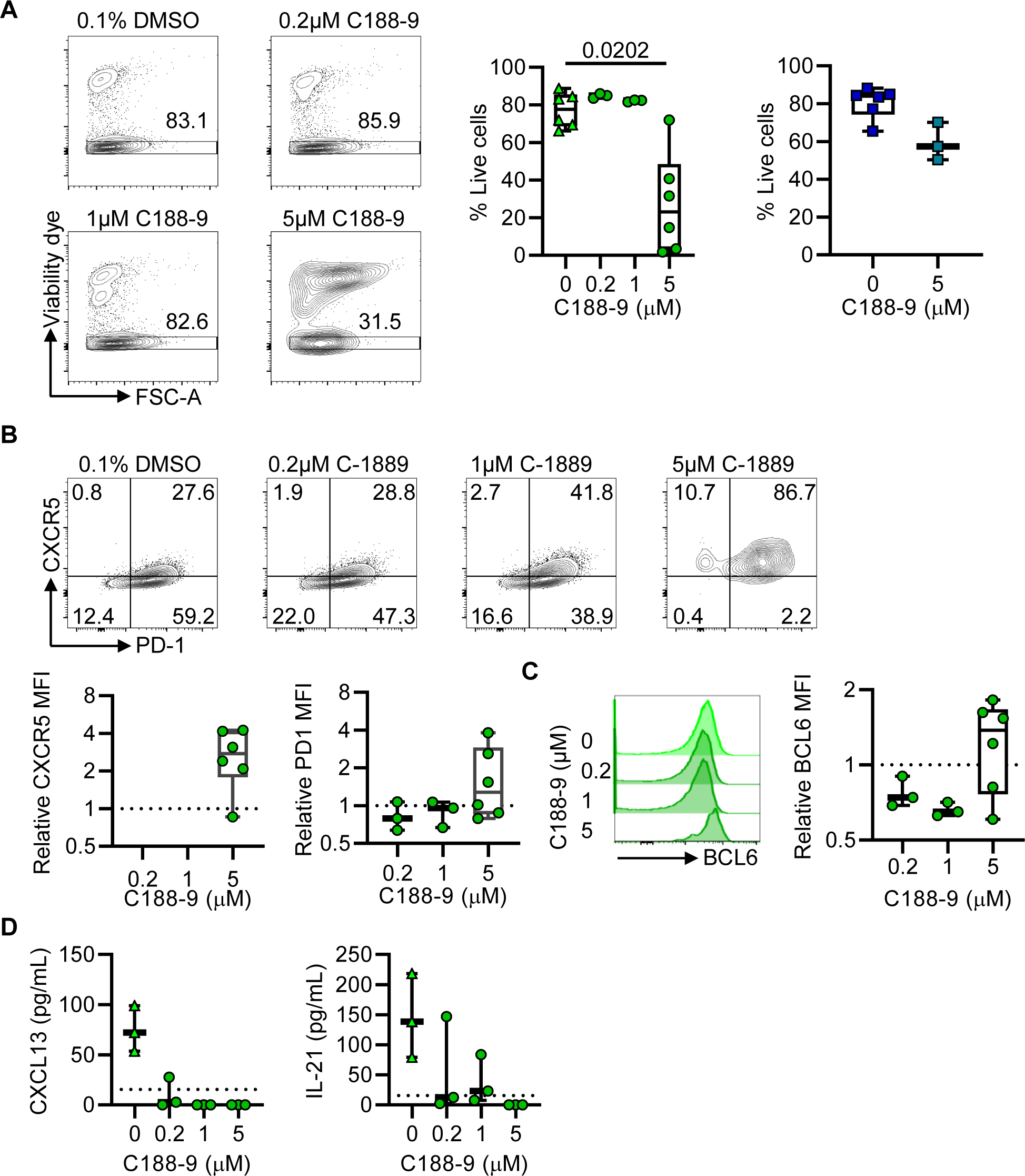
STAT3 signaling is required for *in vitro* TFH viability and functional capacity. pSTAT3 inhibitor C188-9 was dissolved in DMSO. All cell culture samples were adjusted to a final concentration of 0.1% DMSO. **(A)** Cell viability of *in vitro* TFH differentiated in presence of C188-9 or DMSO. Kruskal-Wallis test with Dunn’s multiple comparison test was used. Viability of control CD4 cultured with C-1889 was quantified. **(B)** CXCR5 and PD1 expression by *in vitro* TFH cultured with C188-9, relative to DMSO treated control. **(C)** BCL6 expression by *in vitro* TFH cultured with C188-9, relative to DMSO treated control. **(D)** Cytokine production as quantified by supernatant ELISA for CXCL13 and IL-21. For **(A-C)** 0 and 5 µM n=6, for 0.2 and 1 µM n=3. **(D)** n-3.

This data show that while STAT4 signaling may be redundant for the initial differentiation of TFH, STAT3 signaling is essential for the survival of CD4 T cells in conditions favoring TFH differentiation. In addition, even small changes in STAT3 signaling, while not affecting differentiation, play an important role in promoting function.

### Differentiation of CD4 T cells in TFH favoring conditions upregulates core TFH transcriptional signature

While BCL6 is the master transcriptional regulator of TFH, as with all CD4 T subsets there are a complex network of up and down-regulated genes required to promote both phenotype and function. Whole transcriptome analysis of *in vitro* TFH versus naïve CD4 T cells, tonsillar TFH (CXCR5^hi^PD1^hi^CD25^-^TNFR2^-^) and TFR (CXCR5^hi^PD1^+^CD25^hi^TNFR2^+^) (gating example shown in Fig S2A.) revealed that by principle component analysis *in vitro* TFH formed an independent cluster most closely related to *in vivo* TFR (Fig. 4A). Differential gene analysis confirmed this with *in vitro* TFH having 991 shared genes with tonsil TFR being differentially regulated as compared to naïve CD4s, versus only 295 genes shared with tonsil TFH (Fig. 4B). Analysis of the genes shared by *in vitro* TFH and TFR using GO term biological processes analysis displayed an overrepresentation of genes associated with actively cycling cells (Fig. 4C).

**Figure 4.**
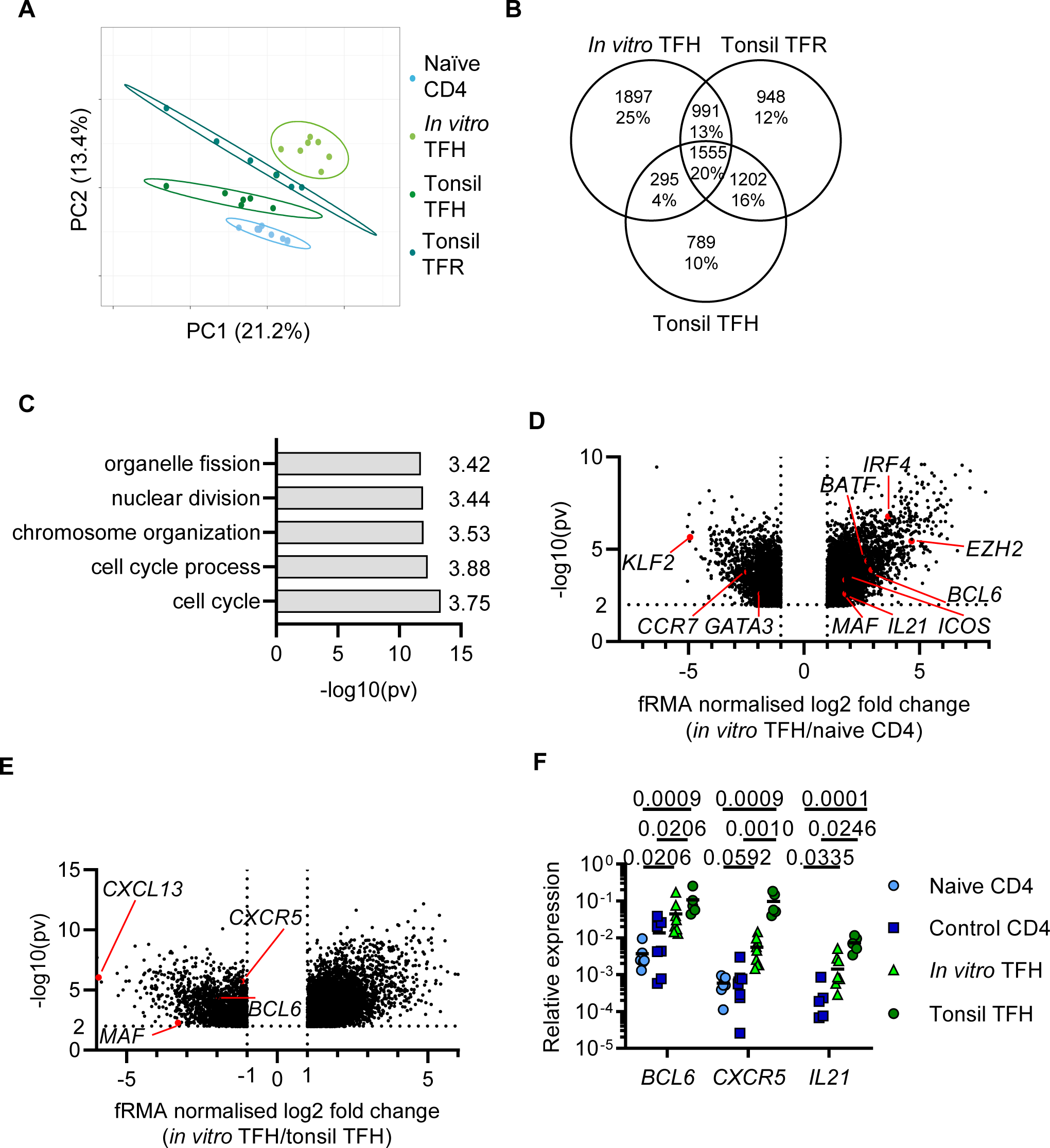
In vitro TFH upregulate the core *in vivo* TFH transcriptome. **(A)** Principal component analysis of whole transcriptome microarray data for indicated cell types. **(B)** Number of genes with a mean minimum two-fold upregulation for indicated cell types and overlap, as compared to naïve CD4. **(C)** GO term biological processes analysis of 1487 genes upregulated by in vitro TFH and tonsil TFR. Fold enrichment is annotated. **(D-E)** Genes with a minimum of 2-fold differential mean expression between naïve CD4 and *in vitro* TFH **(D)** and *in vitro* TFH and tonsil TFH **(E). (F)** Quantification of core TFH lineage factors by qPCR relative to housekeeping genes *18S* and *ACTB*. For each gene, Tukey’s multiple comparison test was used. In samples for which *IL21* transcript was not detected, no value is plotted. For **(A-E)**, naïve CD4 n=8, *in vitro* TFH n=7, tonsil TFH and tonsil TFR n=6. For **(F)**, naïve CD4 n=8, control CD4 n=7, *in vitro* TFH n=9, tonsil TFH n=6.

Importantly however there was limited evidence that *in vitro* TFH conditions resulted in the generation of Tregs, as although limited *FOXP3* transcript upregulation was seen, protein expression was absent (Figure S2B-C). Furthermore, tonsil TFR displayed unique upregulation of hallmark regulatory gene transcripts *CTLA4, PRDM1* and *IKZF2* (Helios) (Figure S2D-F).

Genes shared uniquely between the two tonsil derived populations included both *CXCL13* and *CXCR5*, as well as a downregulation of *S1PR1* as compared to circulating naïve CD4s, demonstrating their tissue resident heritage. Furthermore, the transcription factors *BACH1* and *MYC* were also differentially regulated, demonstrating that *in vitro* TFH reach an intermediate TFH phenotype, with respect to GC TFH. Among the 1555 genes shared by all three populations were transcripts associated with the TFH lineage such as *BCL6*, *ICOS*, *IL21* and the microRNA *MIR155HG*. GO term biological process analysis showed an enrichment for transcripts involved in T cell activation such as “regulation of T cell receptor signaling pathway” (p = 0.0305) and “T cell differentiation involved in an immune response” (p = 0.0388).

While this analysis showed that *in vitro* TFH only shared approximately 25% of the gene changes with tonsil TFH, analysis of the significantly upregulated genes between *in vitro* TFH and naïve CD4 T cells again revealed *BCL6*, *IL21* and *ICOS as well as* many other core TFH associated genes including *MAF, BATF, IRF4, and EZH2,* alongside downregulation of factors associated with naïve CD4 T cells, such as *KLF2* and *CCR7*, and downregulation of the Th2 lineage factor *GATA3* (Fig. 4D). Interestingly however, while many TFH associated features were upregulated after *in vitro* TFH differentiation in comparison to naïve CD4 T cells, some factors remained significantly downregulated when compared to *in vivo* tonsil TFH, including *CXCL13*, *CXCR5* and *BCL6* (Fig. 4E). RT-qPCR analysis of specific genes confirmed these findings, confirming as had been seen at the protein level that *BCL6*, *CXCR5* and *IL21* are upregulated by *in vitro* TFH compared to naïve CD4 T cells or control CD4 T cells, but remain significantly lower than the expression found in tonsil TFH (Fig. 4F). Thus, use of TFH favoring conditions *in vitro* can promote upregulation of many of the core features of TFH by naïve CD4 T cells, but not to the same extent as is observed in those TFH derived *in vivo*.

### IL-2-STAT5 signaling restrains TFH differentiation in vitro

Pathway analysis predicted an upstream activating role for the common γc cytokine IL-15, alongside the growth factor Amphiregulin, in promoting the *in vitro* TFH phenotype (Fig 5A). However, *AREG* transcript was detected at only baseline levels in *in vitro* TFH (Fig. 5B). We explored IL-15 in the context of the γc cytokine family; IL*2* was significantly increased in *in vitro* TFH compared to both naïve CD4 T cells and *in vivo* TFH or TFR, as was another γc cytokine, *IL9.* Of the other γc cytokines *IL4* and *IL7* had a similar pattern to *IL15* while *IL21* as previously shown was highly upregulated in all follicular helper cell populations (Fig. 5C). *In vitro* TFH also showed heightened expression of the IL-2 receptor complex genes (*IL2RA, IL2RB* and *IL2RG*) compared to both naïve CD4 T cells and *in vivo* TFH. Expression of *IL15RA* and *IL21R* was also increased in *in vitro* TFH, while *IL9R* expression was similar in all cell types analyzed and *IL4R* and *IL7R* expression were reduced in *in vitro* TFH compared to naïve CD4 T cells (Fig 5D). While IL-2 can be difficult to detect after release due to its high turnover, IL-2^+^ cells were readily detectable by cytokine capture in both *in vitro* TFH and control CD4 T cells after 6 days of culture (Fig 5E). IL-9 was also easily detectable in the supernatant after *in vitro* TFH culture by ELISA, but IL-15 was poorly detected (Fig. 5F). Upon polyclonal stimulation it was found that unlike naïve CD4 T cells a substantial proportion of *in vitro* TFH produce IL-2, with a subset of IL-2^+^ cells also producing IL-9 (Fig. 5G).

**Figure 5.**
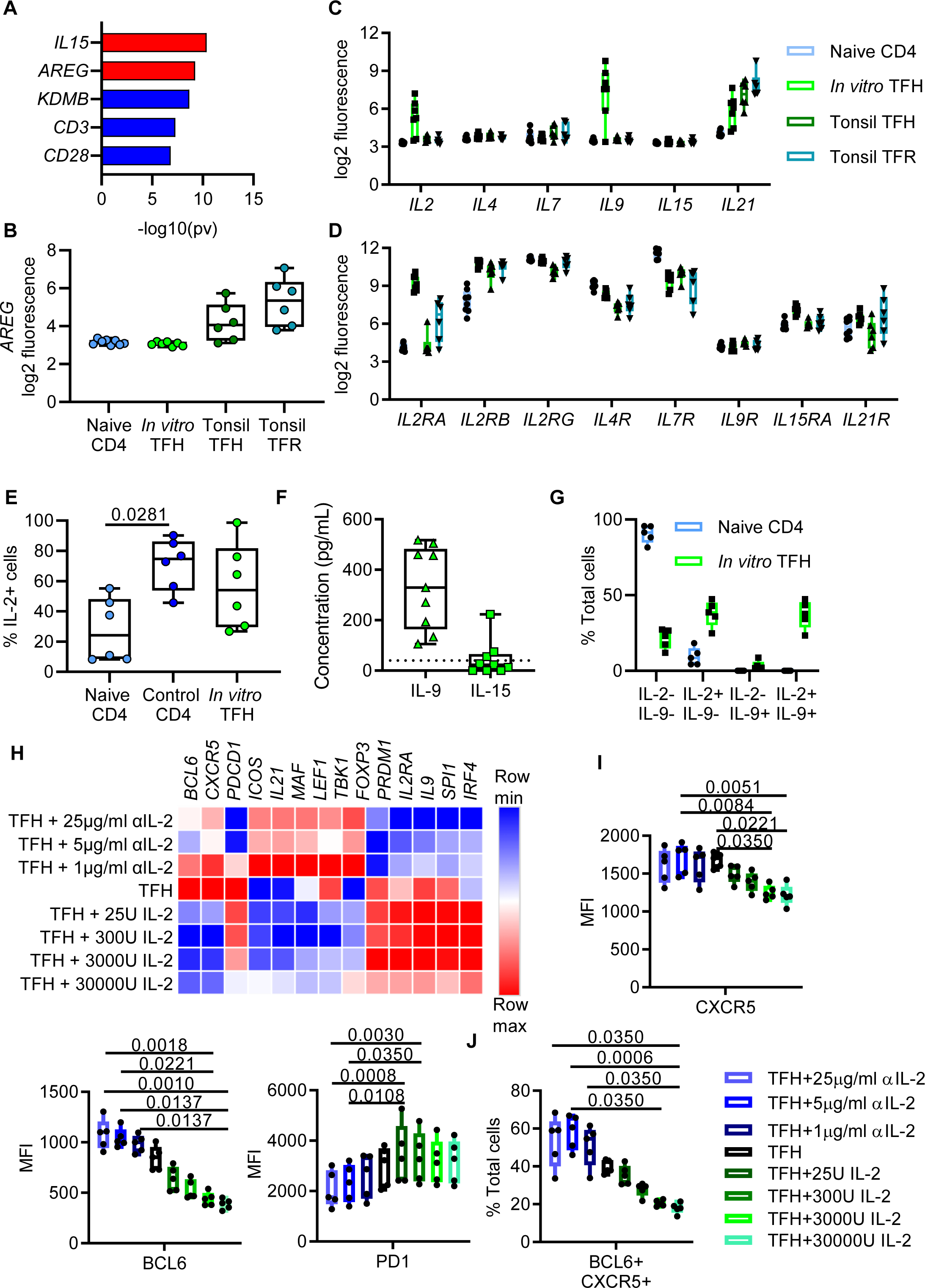
TFH produce IL-2 during differentiation, setting up an autocrine negative feedback loop. **(A)** Ingenuity Pathway upstream regulator analysis was performed on differentially expressed genes between naïve CD4 and in vitro TFH. Regulatory pathways in red are predicted to be activated, pathways in blue are inhibited. **(B)** log2 transformed Amphiregulin transcript expression for indicated cell types. **(C-D)** Common gamma-c cytokine **(C)** and receptor **(D)** family transcript expression by naïve CD4, *in vitro* TFH, tonsil TFH and tonsil TFR was quantified. **(E)** IL-2 surface capture assay was used to quantify IL-2 production by indicated cell types during culture, without polyclonal stimulation. **(F)** IL-9 and IL-15 was quantified from *in vitro* TFH culture supernatants by ELISA. **(G)** Intracellular IL-2 and IL-9 expression following PMA/Ionomycin polyclonal stimulation in the presence of Brefeldin A. by indicated cell types. **(H)** mRNA transcript of *in vitro* TFH cultured in the presence of increasing concentrations of αIL-2 monoclonal antibody or exogenous recombinant IL-2 was quantified via Fluidigm PCR array. Heatmap shows mean transcript expression. **(I-J)** Protein expression of core TFH lineage markers was quantified for *in vitro* TFH, Friedman test with Dunn’s multiple comparison test was used. For **(A-D)**, naïve CD4 n=8, *in vitro* TFH n=7, tonsil TFH and tonsil TFR n=6. For **(E)** n=6. For **(F)** n=9. For **(G-J)** n=5.

Fitting with induction of IL-9 by other immune cell populations, the presence of TGF-β and Activin-A was however vital for secretion of IL-9 from *in vitro* TFH (Fig S3A). Analyzing expression of core T helper 9 transcription factors revealed that *in vitro* TFH upregulate *IRF4*, but not *SPI1, FOXO1* or *STAT6* compared to naïve CD4 T cells suggesting an absence of a TH9 phenotype (Fig S3B). Although IL-9 is produced by *in vitro* TFH, *IL9R* did not appear to be abundant transcriptionally (Fig 5D), nor was it detectable at a protein level (Fig.S3C). Concomitant with this, differentiation of TFH in the presence of a neutralizing anti-IL9 antibody did not affect CXCR5, PD1 or BCL6 expression when compared to isotype treated cells (Fig S3D).

While neutralizing IL-9 appeared to have minimal effects on TFH differentiation and function neutralizing IL-2 during *in vitro* TFH differentiation resulted in significant upregulation of core TFH genes including not only *BCL6*, but also *MAF*, *LEF1* and *TBK1,* with *IL21*, *ICOS* and *CXCR5* (Fig. 5H). In fact, regulation of the genes appeared to tightly correlate with the relative abundance of IL-2 signaling, with increasing doses of recombinant IL-2 results in progressive down regulation of the same transcripts (Fig. 5H). In contrast to this *PRDM1*, *IL9*, *IRF4*, *PDCD1* and *IL2RA* itself were most highly expressed in TFH exposed to high dose IL-2 and lowest when cells were treated with high doses of anti-IL2 antibodies (Fig 5H). This effect was reproducible at the protein level with BCL6 and CXCR5 inversely correlated with the amount of IL-2 available, while PD-1 positively correlated with IL-2 concentrations (Fig 5I). Overall, this resulted in a higher proportion of CXCR5+ PD1+ BCL6+ TFH when IL-2 was depleted during differentiation (Fig. 5J). Collectively this suggests the generation and function of TFH is acutely sensitive to the presence of IL-2 signaling, and excessive IL-2 production after *in vitro* TCR engagement limits acquisition of TFH phenotype.

## Discussion

In this study, we demonstrate a system by which human TFH analogs can be generated and expanded. We show that these *in vitro* derived TFH upregulate BCL6, PD1 and CXCR5, and promote B cell differentiation in co-culture. Moreover, we determined that acquisition of TFH phenotype is hindered in absence of either STAT3 or SMAD2/3 signaling cytokines. In addition, we compared the phenotype of *in vitro* TFH with tonsil derived germinal center TFH and TFR, concluding that *in vitro* derived cells upregulate key TFH differentiation transcription factors; such as BCL6 and MAF, but do so in an intermediate fashion with respect to tonsillar TFH. Finally, we found this intermediate *in vitro* TFH phenotype was a result of their production and response to IL-2.

Cytokines influencing TFH differentiation have been intensely studied, with individual reports showing that IL-6, IL-12, TGF-β, Activin-A and IL-1β (28, 31, 32, 39) all have roles in promoting TFH differentiation. Our work integrates these cytokines alongside ICOS signaling to generate a robust system capable of producing TFH-phenotype cells *in vitro*. These cells were capable of promoting B cell differentiation, likely due to their BCL6-dependent capacity to produce IL-21 and CXCL13 (6, 40, 41). By investigating the importance of individual cytokines (grouped as either STAT3/4 or SMAD2/3) signaling pairs, we demonstrate the relative importance of IL-6 vs IL-12 and TGF-β vs Activin-A for human TFH differentiation. We confirm that IL-12, and to a lesser extent IL-6 (both via STAT3 signaling) govern IL-21 and CXCR5 induction in human TFH (29). IL-12, unlike IL-6, appears necessary for optimal differentiation, with additional loss of IL-6 further reducing acquisition of TFH phenotype, a result that agrees with previous works demonstrating IL-12 as the predominant STAT3 signaling cytokine for human TFH, with IL-6 having a supplementary effect (29, 32). Confirmation of the importance/central role of STAT3 was shown using the small molecule inhibitor C188-9, which blocks STAT3 signaling (38), and here negatively influenced BCL6 expression, and the production of both IL-21 and CXCL13. Interestingly higher concentrations of inhibitor caused widespread cell death in TFH differentiation, but not control T cell conditions. This indicates that sustained STAT3 signaling is a requirement for TFH survival, but less so for other CD4 T cell subsets. As this was not observed when IL-6 and IL-12 were removed, it also implies autocrine production of other STAT3 signaling factors by *in vitro* TFH, mostly likely IL-21 given its known role in TFH differentiation and reduced production in the presence of low doses of STAT3 inhibition.

Activin-A and TGF-β, when combined with IL-12, have both been proposed to be important signaling cytokines for human TFH development (31). Furthermore, TGF-β is able to mediate the chromatin organizer SATB1, which results in increased ICOS expression of activated CD4 T cells, as well as FOXP3 repression, leading to increased PD1 expression, and increased TFH/TFR ratio (42). In agreement with this, our *in vitro* system shows that Activin-A and TGF-β signaling can regulate the *BCL6*/*PRDM1* ratio, a key pivot in determining TFH differentiation and function. Our data however suggest TGF-β has the greater individual role, as removal of Activin A alone has only limited downstream consequences while TGF-β was required for optimal CXCR5 and PD-1 expression. Taken together these experiments show that both STAT3 and SMAD2/3 signaling is required for the generation of phenotypically accurate and functional human TFH.

Transcriptomic comparison of *in vitro* TFH versus naïve CD4 and tonsillar TFH suggests successful recapitulation of the *in vivo* cytokine signals required for early TFH differentiation, including BCL6 upregulation, which promotes TFH fate through pleiotropic action (43) and MAF, another non-redundant promoter of early TFH differentiation (44). Consistent with this acquisition of transcription factor activity, the expected downstream effector proteins (CXCR5, PD1, ICOS, IL21, and CXCL13) are all upregulated, with concomitant downregulation of CCR7. Despite this, *in vitro* TFH retain lower expression of *BCL6* and *MAF*, and many key TFH associated genes, compared to tonsillar TFH. TFH differentiation can be considered as a step-wise process, with initial DC priming resulting in pre-TFH that migrate to the T-B border receive further signals that promote their migration into the B cell zone. GC TFH, which are the predominant type found in human tonsil, meanwhile require interactions with GC B cells for full acquisition of their phenotype (45). It could be speculated therefore that signals produced *in vitro* result in a pre- or intermediate TFH phenotype, with further distinct signals required for further maturation.

Perhaps surprisingly *in vitro* TFH transcriptionally overlap more consistently with tonsil TFR than TFH, and this highlights at least one missing signal that is almost certainly involved in the final maturation of TFH. This overlap of *in vitro* TFH and tonsillar TFR is not a result of *in vitro* cell possessing an immune-suppressive regulatory signature, since TFR upregulated the hallmark genes of functional Tregs, such as *FOXP3*, *CTLA4*, *PRDM1* and *IKZF2* (46–48), but *in vitro* TFH did not. Instead the overlap appears the result of TFR and *in vitro* TFH, but not TFH, sharing common upregulation of cell cycle pathways. One of the primary upstream regulators associated with this is cytokine signaling. We find that *in vitro* TFH express high levels of both the high affinity IL-2 receptor CD25 and IL-2 itself, a potent T cell proliferation factor.

IL-2 has been widely reported as a negative regulator of TFH differentiation in both mouse models and humans; signaling through STAT5 to shift the BCL6/PRDM1 balance away from BCL6 (20, 49, 50). IL-2 production occurs following high-affinity T cell receptor engagement on naïve T cells and thus *In vivo* selective expression of IL-2 during T cell priming is therefore required to promote TFH differentiation. Using murine models, it was recently shown that cells receiving high affinity TCR ligation produce paracrine IL-2 and become TFH, whilst cells receiving this IL-2 signaling are driven towards the Th1 lineage (51). Moreover, other cell types in lymphoid tissue microenvironment help quench IL-2 from the micro-environment, most notably CD25^+^ T regulatory cells and CD25-secreting dendritic cells, further facilitating TFH differentiation (52, 53). The high-affinity antibodies used here to promote TCR signaling *in vitro*, combined with homogenous nature of the naïve CD4 T cell cultures, thus results in a high proportion IL-2 producing cells in the absence of any quenching or competition for IL-2. Reducing IL-2 signaling in this pro-TFH environment results in upregulation of a core TFH transcriptional signature including *BCL6*, *MAF*, *LEF* and *TBK1*, and reduced *PRDM1,* and key functional molecules such as *IL21* and *CXCR5*, limiting IL-2 signaling also results in downregulation of *IL2RA*, driving *in vitro* TFH further towards their *in vivo* counterparts. This highlights the pivotal nature of IL-2 mediated antagonism in regulating terminal TFH differentiation and function.

The *in vitro* differentiation of human TFH will always have limitations in how directly it replicates the *in vivo* environment, however as shown above it does allow dissection of mechanisms that contribute to the differentiation or function of human TFH. For instance, in our *in vitro* setting TFH produced IL-9 in a TGF-β and Activin-A dependent manner, which was further regulated by IL-2 signaling. This IL-9 production did not occur as a result of TH9 phenotype acquisition, a nor does IL-9 act in an autocrine fashion on TFH themselves, unlike IL-2. GC B cells however do express IL-9R (54), and IL-9 signaling from TFH to GC B cells has been reported to promote the formation and exit of memory B cells from the GC in mice (55), whilst a second study shows that IL-9 signaling is required for normal recall responses; with memory B cells expressing IL-9R after primary immunization (56). The *in vitro* TFH system described here could be utilized to examine the role of IL-9 in T dependent regulation of B cells in humans, and also explore how human TFH can be modulated to produce other effector cytokines such as IL-13+ TFH reported to be important in IgE responses or IL-17+ TFH found in the mucosa (37, 57).

In summary, we demonstrate a reliable method of generating cells that closely resemble human TFH. We show that the intermediate stage of TFH differentiation represented by *in vitro* TFH occurs partially as a result of a failure, *in vitro*, to maintain IL-2 homeostasis and that correction of this results in further maturation of TFH. These data highlight the importance of STAT3 cytokines in promoting TFH differentiation, and the ability of STAT5 cytokines to regulate differentiation, while providing a novel system to allow in depth dissection of these processes.

## Methods

### Naïve T cell and B cell isolation by negative selection

15mls of Lymphoprep (Stemcell Technologies) was layered into a 50ml tube, with peripheral blood of healthy volunteers diluted 1:1 in FACS buffer (PBS with 2% fetal bovine serum (FBS) and 1mM EDTA) layered on top. Tubes were centrifuged (400g 40 minutes with no brake). Buffy coat was removed, cells counted and EasySep human naïve CD4 or naïve B cell isolation kits (Stemcell Technologies) were used following manufacturer’s instructions.

### Pediatric tonsil collection and tissue processing

Tonsil samples used were collected under the research project “Investigating the role of the IL-6 cytokine family and gp130 in the maintenance of germinal center integrity” (Imperial College Tissue bank no. R17019), with patient recruitment and consenting being performed by the specialty registrar performing the surgery. Tonsils were processed within one hour of tonsillectomy. The entire tonsil was cut finely with a scalpel and filtered through a 70 µM nylon filter with warm RPMI (ThermoFisher Scientific). The resulting cell suspension was resuspended in (Ammonium-Chloride-Potassium) Red cell lysis buffer for 5 minutes, before being washed twice in RPMI. Tonsil samples were cryopreserved in 90% FCS, 10% DMSO, frozen over liquid nitrogen and stored long term at -80°C. Cells were defrosted and rested in warm RPMI for 3 hours before flow cytometry sorting.

### In vitro TFH culture and B cell coculture

For TFH differentiation wells, 100 µL of 5 µg/mL B7-H2-Fc (ICOS-L-Fc) (R&D systems) was coated onto round bottom 96 well tissue culture plates overnight. Plates were washed twice with PBS before use. Naïve CD4 T cells were re-suspended in AIM-V serum free media (Gibco) at 5×10^5^ cells/mL, supplemented with 10 µL/mL of αCD3/αCD28 Dynalbeads (Invitrogen). 5×10^4^ cells were plated into each well.

After 24 hours, recombinant cytokines were added, with TFH differentiation samples using final concentrations of: 25 ng/mL IL-6, 1 ng/mL IL-12, 10 ng/mL IL-1β, 5 ng/mL TGF-β (all R&D Systems) and 100 ng/mL Activin A (Peprotech). For non-TFH control samples, cells were cultured without ICOS-L-Fc and recombinant IL-7 (Peprotech) was added to a final concentration of 4 ng/mL instead.

In TFH/B cell co-culture experiments, TFH were differentiated as above, and re-suspended at 5 x 10^5^ cells/ml. 5×10^4^ heterologous B cells were plated per well and then cultured with either 100ng/mL IL-21 (R&D Systems) and 5μg/mL αCD40 (Invitrogen) or 2.5 x 10^4^ *in vitro* TFH. After 72 hours, B cell phenotype was analyzed by flow cytometry.

In experiments using the small molecule inhibitors (±)-Lisofylline and XIII C188-9 (Merck), these were added one hour before recombinant cytokines. C188-9 was dissolved in DMSO, so all samples using this inhibitor were adjusted to a final DMSO concentration of 0.1%.

### Flow cytometry and sorting

Viability staining was achieved using either LIVE/DEAD™ Fixable Blue Dead Cell Stain Kit or eBioscience™ Fixable Viability Dye eFluor™ 780 (ThermoFisher), for 30 minutes at 4°C. Surface staining was carried out for 30 minutes at 4°C, followed by 2 FACS buffer washes. Samples were fixed with the Foxp3 Staining Buffer Set (ThermoFisher Scientific) for 30 minutes at 4°C. After fixation, cells were washed with FACS buffer, then washed with 150 µL of permeabilization buffer (Foxp3 Staining Buffer Set). Antibodies were diluted in permeabilization buffer, added to cells, and incubated for 60 minutes at 4°C. Cells were washed twice with FACS buffer and acquired with either a BD LSRFortessa or BD Symphony (BD Bioscience).

Cells used for Fluorescence-activated cell sorting (FACS) experiments were stained with surface antibodies for 30 minutes at 4°C, and stained with 0.1 µg/mL 4′,6-diamidino-2-phenylindole (DAPI) for 10 minutes at room temperature immediately before sorting. Flow cytometry data was analyzed with Flowjo V10 (Treestar).

### RNA extraction and cDNA conversion

Cells were resuspended in RLT buffer (Qiagen) supplemented with 0.1% β-mercaptoethanol, and stored at -80°C until extraction. RNA was extracted using the Qiagen RNeasy mini-kit, following manufacturer’s instructions, to maximize RNA concentration in a 30 µL volume of RNAse free water. For samples used in single transcript assays, RNA concentration and quality were quantified using a NanoDrop1000 (Thermofisher). For samples used in microarrays, RNA concentration and RIN (RNA Integrity Number) was quantified using a Tapestation2200 (Agilent technologies). For all microarray samples, a minimum RIN value of 8.0 was achieved. For samples used in single transcript assays, cDNA was converted using GoScript™ Reverse Transcriptase (Promega). For samples used for microarray analysis, Superscript3 III First-Strand Synthesis System (Thermofisher) was used, all as per manufacturer’s instructions.

### qPCR and Fluidigm array

For all qPCR experiments unless otherwise noted, TaqMan probes and TaqMan Fast Advanced Master Mix (both Thermofisher) were used, with a 384 well plate-Viia7 Real Time PCR system (Applied Biosciences) being used for transcript quantification. For IL-2 / αIL-2 differentiation studies, the 96.96 Dynamic Array IFC (Fluidigm) was used, which included a pre-amplification step, using the TaqMan™ PreAmp Master Mix Kit (Thermofisher). Automated assay mixing was completed using the Fluidigm IFC controller system. After mixing, the plate was analyzed using the BioMark analyzer (Fluidigm), all as per manufacturer’s instructions.

### Whole transcriptome microarray and gene expression analysis

Whole transcriptome analysis was performed using the Affymetrix GeneChip™ Human Genome U133 Plus 2.0 Array. Resulting data was pre-processed using frozen RMA (Robust multiarray analysis). Raw luminescence data generated by the microarray was log2 transformed before downstream analysis. R studio was used to write R scripts used to handle data. Heat maps were generated using the Morpheus tool (https://software.broadinstitute.org/morpheus/). Hierarchical clustering was performed using the One minus Pearson correlation. Venn diagrams were generated using the Venny 2.1 tool (https://bioinfogp.cnb.csic.es/tools/venny/). PCA plots were generated using ClustVis (https://biit.cs.ut.ee/clustvis/). PCA plots were generated using singular value decomposition (SVD) with imputation. Gene Ontology (GO) term analysis was performed using the biological processes analysis tool (http://geneontology.org/). Ingenuity Pathways Analysis (IPA) upstream regulator analysis tool was used for comparative analysis of groups of microarray samples in order to identify upstream regulators mediated *in vitro* TFH differentiation.

### IL-2 capture assay and cytokine re-stimulation assays

Cells were stained with IL-2 capture reagent (Miltenyi Biotec) as per manufacturer’s instructions. Cells were stained for flow cytometry as above, with the APC-conjugated IL-2 detection reagent being included in the staining mix. For cytokine restimulation assays, cells were resuspended in a stimulating media mix containing 50 ng/mL phorbol myristate acetate (PMA) (Sigma-Aldrich) and 500 ng/mL Ionomycin (Emdchemicals) in the presence of 10 µg/mL Brefeldin A (Sigma-Aldrich) for 4 hours at 37°C 5% C0_2_. As an unstimulated control, cells were stimulated for 4 hours with 10 µg/mL Brefeldin A only. Cells were then immediately stained for flow cytometry, with anti-cytokine antibodies included in the intracellular mix.

### Cytokine ELISAs

IL-9 and IL-15 cytokine ELISAs (Invitrogen) were performed following manufacturer’s instructions, using half area flat bottom 96 well plates. Supernatant samples were defrosted and 35 µL was added to wells. ELISA plates were read, and standard curves calculated using a SpectraMax i3x (Molecular Devices).

### In vitro IL-2 stimulation and blockade of IL-2 and IL-9

In experiments exploring the role of IL-2 in TFH differentiation, naïve CD4 T cells were isolated as previously described before being resuspended in media containing a final concentration of either recombinant human IL-2 (Peprotech), LEAF™ Purified anti-human MQ1-17H12 IL-2 Antibody (BioLegend), or 5 µg/mL of rat IgG2a isotype control antibody (BioLegend), as indicated prior to *in vitro* TFH differentiation. For experiments blocking IL-9 signaling, a final concentration of 5 µg/mL purified anti-human MH9A3 IL-9 Antibody (BioLegend), or 5 µg/mL of rat IgG1k isotype control antibody (BioLegend) was used instead.

## Resource Identification Initiative

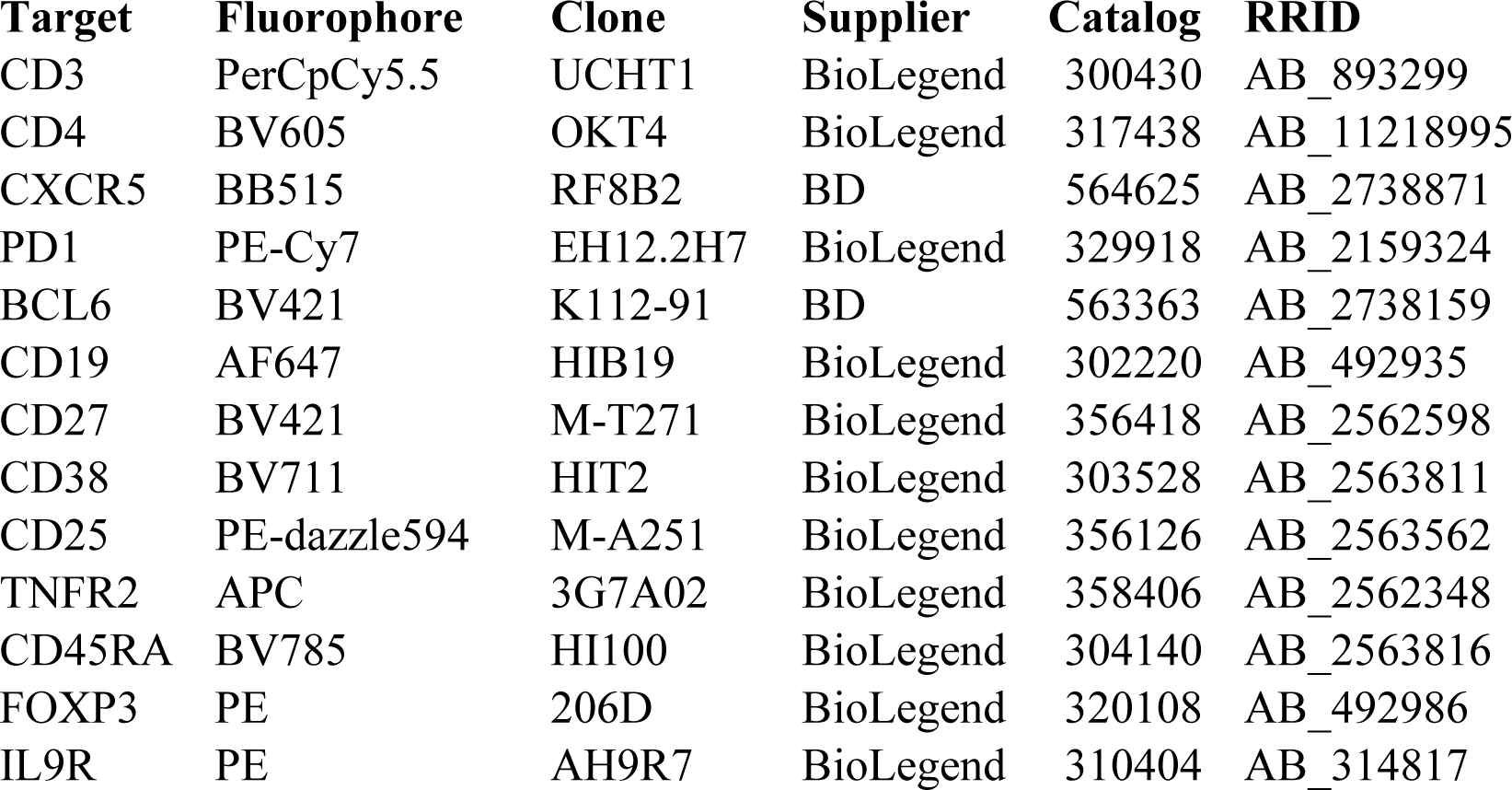

## Author Contributions

WSF, LL-I, and JAH designed experiments. WSF and LL-I carried out and analyzed the experimental work. WSF and JAH wrote the manuscript. LL-I, GSW, MJR, GC, CML, and JAH provided feedback.

## Data Availability Statement

The data that support the findings of this study are available from the corresponding author upon reasonable request.

The microarray datasets generated for this study can be found in the GEO repository “GSE191251 Whole transcriptome analysis of human CD4 T cell subsets”. Private access tokens can be supplied for peer review as required.

---

**Supplementary figure 1.**
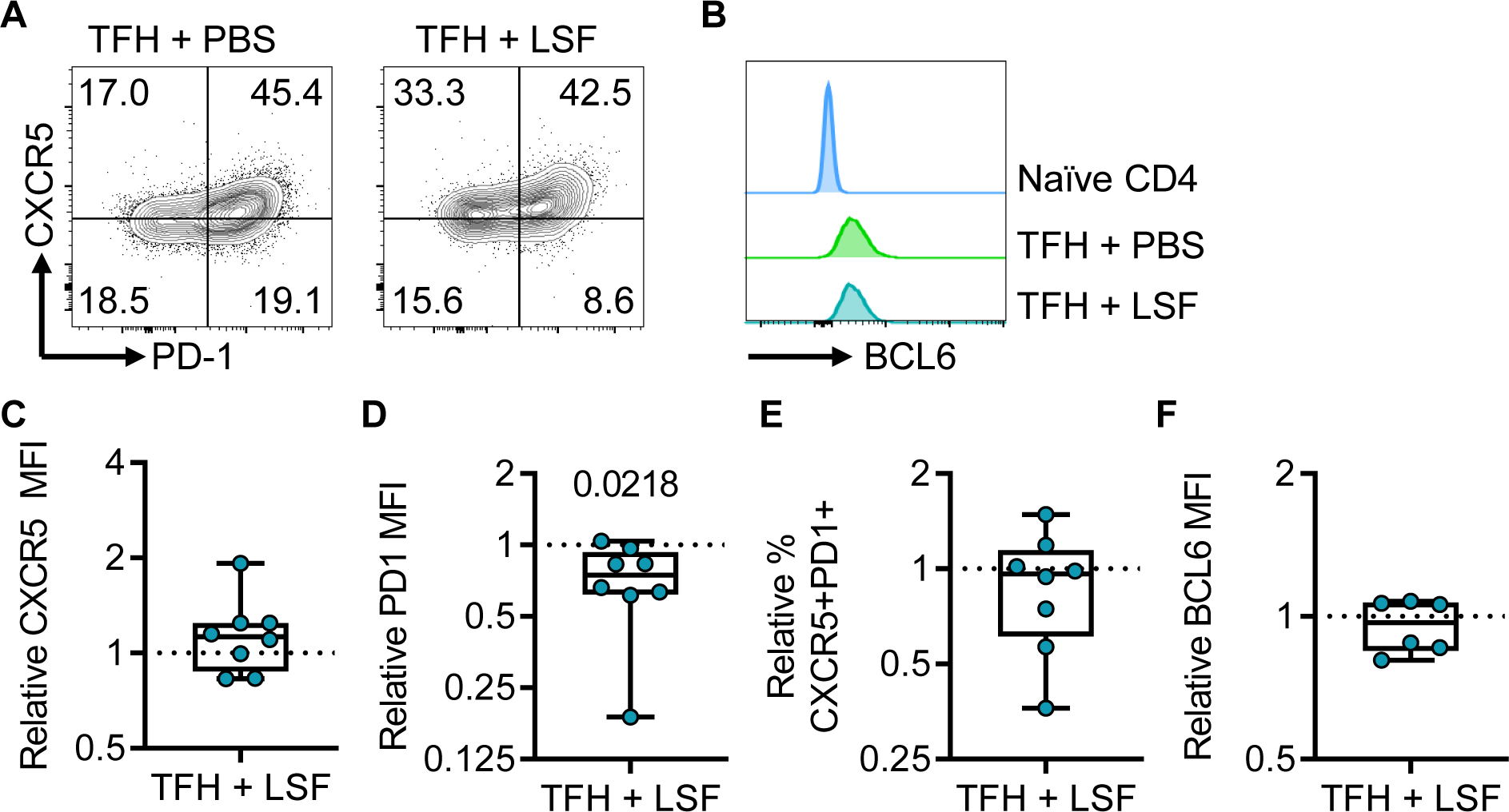
(±)-Lisofylline inhibits PD1 upregulation by TFH. **(A)** Representative plots of PD-1 and CXCR5 expression for in vitro TFH cultured as in Figure 1, alongside *in vitro* TFH cultured in presence of 100μM (±)-Lisofylline (LSF). Numbers represent percent of cells in each gate. **(B)** Median representative histograms for BCL6 expression. **(A-B)** pre-gated on live, single, CD4+ cells. **(C-F)** Change in MFI for TFH cultured in the presence of LSF as compared to paired TFH cultured without LSF for CXCR5 MFI **(C)**, PD-1 MFI **(D)**, TFH conversion (% CXCR5+ PD1+ cells) **(E)** and **(F)** BCL6 MFI. For **(C-F)** the Wilcoxon signed paired ranked test was used. For **(C-E)** n=8, for **(F)**, n=6. For **(A-E)** n=8 biological replicates, across 3 independent experiments. For **(F)** n=6 biological replicates pooled from 2 independent experiments. RM one way ANOVA with Tukey’s multiple comparison test was used. Box and whisker plots show IQR and median with min/max points.

**Supplementary Figure 2.**
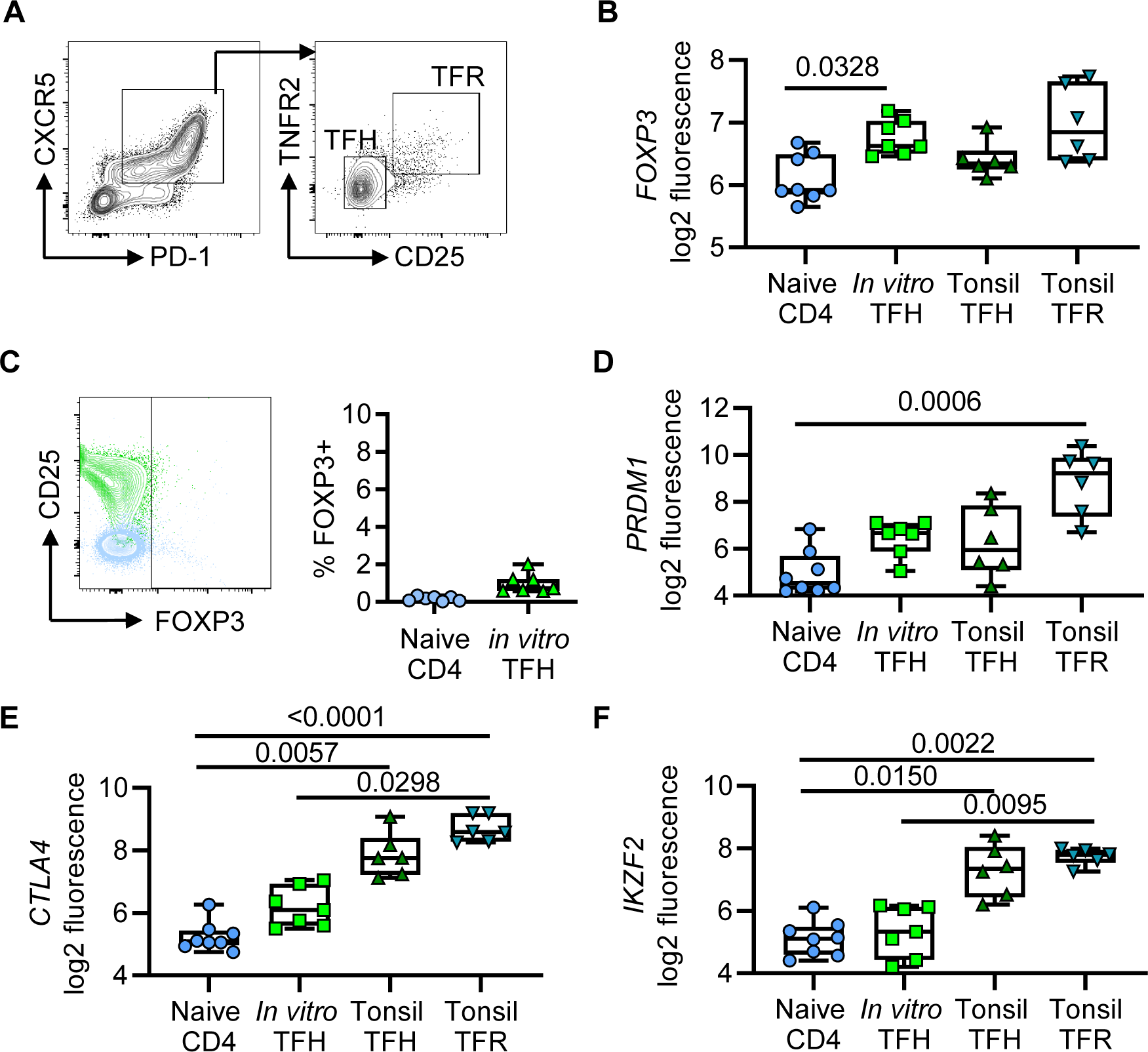
*In vitro* TFH do not express Foxp3 or regulatory transcripts. **(A)** Gating strategy used to sort tonsillar TFH and TFR cells. Plots are pre-gated on live, single, CD3+ CD4+ CD45RA-lymphocytes. **(B)** log2 transformed *FOXP3* transcript expression for indicated cell types. **(C)** Quantification of FOXP3 protein expression on naïve CD4 (negative control) and *in vitro* TFH. **(D-F)** log2 transformed transcript expression of *PRDM1*, *CTLA4* and *IKZF2*. For **(B,D-F)** Kruskal-Wallis test with Dunn’s multiple comparison test was used. Naïve CD4 n=8, *in vitro* TFH n=7, tonsil TFH and tonsil TFR n=6 biological replicates, pooled from 3 independent *in vitro* experiments. Box and whisker plots show IQR and median with min/max points.

**Supplementary Figure 3.**
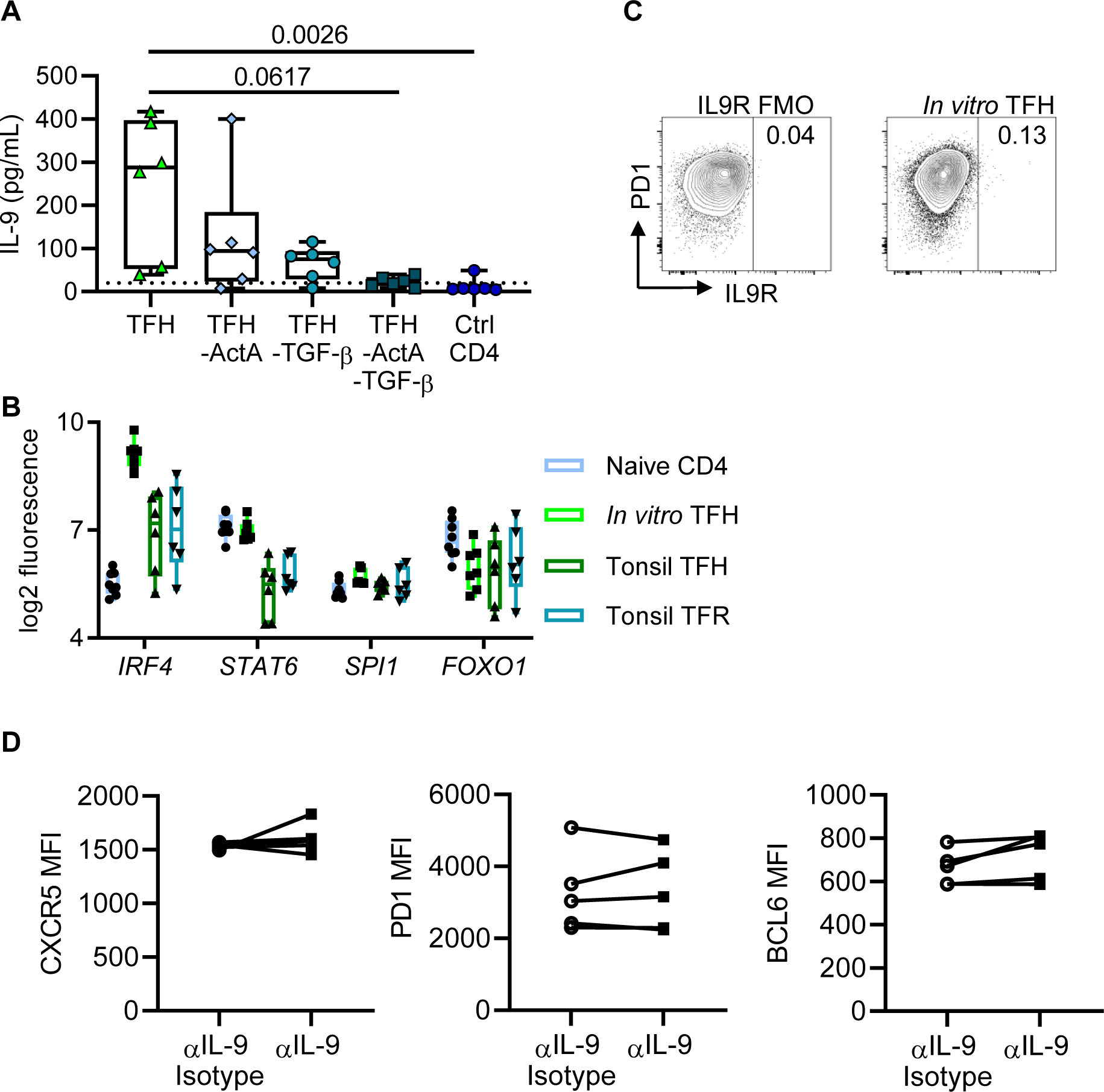
*In vitro* TFH are not IL-9 cells, and are IL-9 insensitive. **(A)** IL-9 was quantified from *in vitro* TFH culture supernatants with indicated cytokine removals by ELISA. Friedman test with Dunn’s multiple comparison test was used. **(B)** log2 transformed transcript expression of Th9 associated factors for indicated cell types. **(C)** IL9R receptor was stained for by flow cytometry, representative plot shows *in vitro* TFH alongside *in vitro* TFH IL9R fluorescence minus one (FMO) negative control. **(D)** Protein expression of core TFH lineage markers was quantified for *in vitro* TFH cultured in presence of anti-IL-9 neutralising antibody or isotype control. For **(A)** n=6 biological replicates across 2 independent experiments. For **(B)** naïve CD4 n=8, *in vitro* TFH n=7, tonsil TFH and tonsil TFR n=6 biological replicates across 3 independent *in vitro* experiments. Box and whisker plots show IQR and median with min/max points. For **(D)** n=5 biological replicates across 2 independent experiments.

